# Effects of High Copper Exposure on Development and Survival During Early Ontogenesis in the Common Toad

**DOI:** 10.1101/471466

**Authors:** ElenaV. Dmitrieva

**Affiliations:** Lomonosov Moscow State University, Faculty of Biology, Moscow, Russia, 119234

## Abstract

Exposure to environmental pollutants contributes to the global decline of amphibians. Amphibian embryos are unique models for studying the effects of various toxic agents on developmental processes. Some of the most detrimental toxic agents are heavy metals, the study of which is a priority in ecotoxicology. Copper is a microelement required in many biological systems, but it can be toxic at concentrations higher than normal physiological range. The purpose of the present study was to analyze the toxic effect of high concentrations of Cu (10, 25, and 50 mg/L) on Bufo bufo embryos. The present study shows that short-term (48 h) toxicity test can fail to register toxicological effect of Cu, whereas long-term toxicity test demonstrates that all analyzed concentrations are lethal to common toad embryos. Early stages of development (stages 4-8, Gosner 1960) are shown to be rather tolerant to copper, even at high concentrations. Effects of copper begin to manifest themselves at the blastula stage (stages 8-9, Gosner, 1960). The present study also shows that initial egg density affects copper-related mortality rate of embryos. If egg density is high, copper-related mortality rate is sometimes decreased, possibly because jelly coats of eggs bind great amounts of dissolved copper. It is also shown that clutches can differ in sensitivity to different copper concentrations. Taken together, our data demonstrate that the conditions of the experiment (such as the initial egg density) and individual sensitivity of clutches to the toxic agent considerably contribute to the results of the ecotoxicological experiment.

---

Dramatic decline of amphibian populations, including local mass extinctions, has been evident worldwide since the 1980s (Darevsky, Orlov, ‘88; Stuart et al., 2004; Reading, 2007). Amphibians are recognized as the most threatened class of vertebrates on the planet (Stuart et al., 2004). The decrease in the number of amphibian species is one of the principal threats to global biodiversity. It is believed that this decrease is caused by destruction and transformation of environments, pollution, pesticide usage, introduced species, climate change, diseases, and rising levels of UV-B radiation (Blaustein, Kiesecker, 2002; Collins, Storfer, 2003; Alford, 2010; Fonovich et al., 2016; Weir, 2016ab). Many cases of amphibian populations suffering from chemical pollution of environments with various pesticides (Cooke, 1973; Brooks, 1981; Alvarez, 1995; Davidson, 2004) and heavy metals (Linder, Grillitsch, 2000; Fedorenkova et al., 2012; Flynn et al 2015; Fonovich et al., 2016; Weir, 2016ab) are known. It is believed that the main cause of decreasing amphibian abundance is the vulnerability of early stages of development (Carey, Bryant, 1995; Mann et. al., 2009).

Industrial enterprises are the main sources of chemical pollution of waterbodies. This is especially true of chemical and petrochemical plants, oil refineries, productions of new synthetic materials, pesticides, and detergents, and plants for thermal processing of solid and liquid fuels. Some of the most detrimental toxicants are the heavy metals cadmium, copper, lead, zinc, and others, considered a priority in ecotoxicological studies (Isidorov, 1997), but the effects of metals and metalloids on amphibians understudied (Hopkins, Rowe, 2010; Flinn et al., 2015). Heavy metals are among the most persistent of pollutants in the ecosystem such as water, sediments and biota because of their resistance to decomposition in natural condition. Under certain environmental conditions, heavy metals might accumulate up to toxic concentrations levels, and cause ecological damage (Bai et al., 2011). Heavy metals can accumulate in both eggs and tadpoles of various amphibian species, affecting their development (Puky, Oertel, 1991; Herkovits, Pérez-Coll, 2007; Fonovich et al., 2016). Copper and lead are among the most common metal contaminants, found in high concentration in many areas, especially those affected by urban and industrial discharges or mining activities (Fatoki, Mathabatha, 2001; SFEI 2015). In recent years, concentrations of copper in the environment have increased dramatically due to widespread use such as electrical generator, radio and television sets, heating system water pipes and wiring (Johnson et al., 2007). In a recent ecological risk assessment of 24 chemical stressors on amphibians, Fedorenkova et al. (2012) ranked Cu exposure as the second most important risk in the Netherlands.

Copper is a microelement required for the functioning of many biological systems (Leary et al., 2009). It is contained in cytochrome oxidase, hemocyanin, some hormones, etc. It is a key element of many intracellular and metabolic processes, such as the synthesis of elastin-related enzymes, collagen, and melanin, and processes providing for the integrity of the central nervous system (O’Dell, 1990). As copper levels in the environment reach some critical level, it triggers the detoxication system of the organism (Herkovits, Helguero, 1998). Elevated Cu levels can increase embryonic and larvae mortality (Lance et al., 2012, 2013; Xia et al., 2012; Flynn et al., 2015), also cause sublethal effects including embryonic deformities (Chen et al., 2007), impaired predator avoidance (García-Munoz et al., 2009; Rumrill, et al., 2016) and other. The boundary at which copper transforms from an activator of biological processes into a toxicant is blurred not only because of the adaptation capacities of the organism, but also because of the process of development as such. The vulnerability of the embryo to copper is different at different stages, because it requires different amounts of this microelement (Dregolskaya, 1993).

Amphibian embryos, used as unique models for studies of damage caused by various toxicants, vividly reveal the consequences of subcellular processes triggered by exposure to the toxicant. As the embryo develops, these hidden changes quickly manifest themselves as malformations, changes in the rate of embryogenesis or termination of embryogenesis, or mortality (Chen et al., 2007; Xia et al., 2012; Lance et al., 2012, 2013; Flynn et al., 2015; Fonovich et al., 2016). Examination of the influence of high concentrations of copper ions on the early development of amphibians can help elucidate its acute toxic effects and estimate the resistance of early stages of embryonic development to such effects. The common toad (*Bufo bufo*) is rather sensitive to copper. For instance, it has been shown in tests of the effects of copper sulfate on tadpoles of different amphibian species that *B. bufo* is more sensitive to copper, compared to *Pelobates cultripes* or *Pelophylax perezi* (Garcıa-Munoz et al., 2010). Therefore, the purpose of the present study was to examine the toxic effects of copper, in particular in high concentrations, on the early development of *B. bufo*.

## MATERIALS AND METHODS

### Study species

The common toad is a species widespread from Western Europe to Eastern Siberia. The abundance of this species in its habitats is rather high; hence, its advantages as a model include accessibility and easiness of collecting. Embryonic and larval stages of development of the common toad are less affected by predator pressure than various frog species (Manteifel, Reshetnikov, 2001); therefore, this factor can be ignored in estimations of mortality in nature. Embryonic and larval stages of the common toad are easy to observe both in nature and in the laboratory. The large clutch size makes it possible to obtain large amounts of genetically uniform materials for various experiments. Each clutch of eggs is a cord up to 10 m long. Females lay eggs in strings containing 1200 to 7200 eggs arranged helically in two rows. These strings twist around underwater objects or lie on the bottom. Laying the eggs, the female usually strains each egg string between stems of water plants or various other objects. Embryonic development depends on temperature and takes 2 to 20 days (Kuzmin, 2012).

Early developmental stages respond to toxicants (Amicarelli et al., 2001; Mandrillon, Saglio, 2007; Bernabo et al., 2008, 2013), including copper ions (Garcıa-Munoz et al., 2010). This species is an indicator of structural changes in anuran assemblages in urbanized environments, because it is a typical forest species capable of successfully reproducing only in forest parks zones of urban areas, intolerant to increased density of the litter, to changes in plant communities, and to changes in microclimate that accompany changes in plant communities (Vershinin, Toporkova, 1981). Therefore, it is one of the first to disappear in anthropogenically transformed ecosystems (Vershinin, 2008).

### Study site

Eggs for the experiments were obtained in a laboratory from breeder toads collected during mass spawning of the common toad in Sterlyazhy Pond within the territory of the Skadovsky Zvenigorod Biological Station, Lomonosov Moscow State University, situated on the right-hand bank of the Moskva River 12 km upstream from Zvenigorod, Odintsovo District, Moscow Oblast, Russia. The pond has artificial origin: it was constructed by damming a small forest stream. The area of the pond is approximately 850 m2; average depth, approximately 1 m; maximum depth, 1.8 m. The bottom is silty-sandy, partly overgrown with vegetation. The water is of medium transparence; the level of copper ions is no higher than the maximum permissible concentration (Cu concentration < 0.001 mg/L).

The pairs were captured in amplexus and brought to the laboratory in approximately 3 h. In the laboratory the pairs were placed in separate aquaria with clean water (Cu concentration < 0.01 mg/L). The captured toad pairs were provided with optimal conditions for egg laying, as close to natural as possible. Each experiment was performed on embryos from one clutch laid by one pair of breeders, i.e., full siblings, which considerably decreased the contribution of the genetic component to variation. Immediately after completion of egg-laying the eggs were distributed over experimental containers. Quality parts of each clutch (not rumpled or deformed) were selected, considerably decreasing the random contribution of dying damaged embryos or unfertilized eggs to the resulting mortality. Stages of development were assessed using Gosner’s normal development Tables (Gosner, 1960).

Egg strings were cut into fragments needed for the experiments, containing 30 or 120 eggs. The choice of the initial density of placement of embryos was determined by earlier studies, which have shown that those populations from which material is taken can be polymorphic in embryo survival rates, depending on initial density (Dmitrieva, 2007, 2013). This effect manifests itself in the fact that approximately 30% of all studied clutches of the common toad demonstrate nonmonotonic dependence of survival rate on initial egg density within the range of 30 to 120 eggs per standard aquarium (∅ 70 mm, h = 30 mm, V = 130 mL); the second maximum of embryo survival rate emerges at a density of 120 eggs per standard aquarium. If experiments are performed at densities higher than 120 eggs, embryo mortality from increased density can be so high as to hide the possible effect of Cu. The density of 30 eggs per standard aquarium is optimal (corresponding to minimum mortality and high synchronicity of development of individual embryos) (Dmitrieva, 2007, 2013). Therefore, the protocols of the experiments take into account the peculiar features of the populations from which material is taken, which makes it possible to obtain most comprehensive data on the influence of the factor in question on embryonic development in the common toad.

### Test solutions

I used in the experiments a standard mix of synthetic dilution water for toxicity tests with freshwater organisms (USEPA, 2002): 48 mg/L NaHCO2, 30 mg/L CaSO4, 30 mg/L MgSO4, and 2 mg/L KCl added to distilled water (pH 7.5–7.8; hardness 40–48 mg/L CaCO3; alkalinity 30–35 mg/L CaCO3). The needed concentration of copper ions was obtained by dissolving the appropriate amount of CuSO4 × 5H2O. Media with Cu concentrations 10 mg/L, 25 mg/L, and 50 mg/L were prepared.

The experiments were performed in 35 mm high glass containers with bottom diameter 70 mm (water volume 130 mL). The corresponding volume of water is sufficient for normal development of common toad embryos (Dmitrieva, 2005, 2007). Water was not added or replaced during the experiment. Dead embryos were not removed, since their removal increases rather than decreases embryo mortality (Dmitrieva, 2005, 2007, 2013). Water is often replaced and/or dead individuals often removed during experiments on embryonic or larval stages of development (American Society for Testing and Materials, 1991; Herkovits, Pérez-Coll, 2003; Garcıa-Munoz et al., 2010; Bernabo et al., 2013 and others). If dead tadpoles or embryos that develop independently from each other (from separate eggs that have no common coat) are removed, this interference can have no considerable effect on the results of the experiment, because it is accompanied only by stirring of the water. However, if dead embryos that develop in a single egg cord are removed, the experimenter has to compromise the wholeness of the structure of the clutch by damaging the common protective coat. I have shown in my earlier experiments that such actions at embryonic stages of development result in increased mortality (Dmitrieva, 2005). Even simple mixing of the water can affect the results of an experiment on embryonic stages of development (Dmitrieva, 2005).

Each experiment was performed in four replications of 30 eggs and three or four replications of 120 eggs (depending on clutch size) from one clutch. Egg cords that contained 30 or 120 eggs were placed in standard aquariums (V = 130 mL, 70 mm ∅). All experiments on embryos were performed the moment of the end of egg-laying, which was taken as 0 time (stage 2–4).

A total of five egg clutches of the common toad have been used in the present study. Three clutches developed in a thermostated room at temperatures within a range of 10–13°C (trial 1: clutches 1–3). Two other clutches were placed in a Binder KB 53 thermostat and developed at a constant temperature of 15°C (trial 2: clutches 4–5), optimal for the development of common toad embryos (Dmitrieva, 2014). The number of dead embryos and stage of development were recorded every 8 h. The experiment ended when all embryos in the control group (i.e., without copper) reached the hatching stage (stage 19). The development time was approximately 230 hours in trial 1 and 130 hours in trial 2. I analyzed treatment effects on percent survival at short-term (48 and 96 h) toxicity tests and to hatching stage.

### Statistical analysis

The results of the experiments were processed using the Statistica 6.0 package (StatSoft Company, http://www.statsoft.ru) (Khalafyan, 2009). The statistical significance of differences between the obtained values of mortality in the experiments in different clutches of the same experiments at different Cu concentrations were estimated using the nonparametric Kruskal–Wallis H test and then use the Mann–Whitney U-test (the nonparametric counterpart of Student’s t-test) as a means to look for differences when the Kruskal–Wallis test is significant. Non-parametric tests were used in connection with a small number of replications of the experiment from one clutch (4 replications of 30 eggs and three and 3-4 replications of 120 eggs). Differences were considered significant at p < 0.05.

The effects of different factors on mortality and rates of development were estimated using the factorial ANOVA. I used a generalized linear mixed effects model to investigate the effects of Cu concentration, clutches and density on mortality of embryos. Mortality was used as the independent variable; copper concentration, clutches and density as the categorical variables. In addition, time of development (i.e. hour since start of experiment) was used as a categorical variable in the long-term test to estimate the influence of the duration of exposure to Cu on mortality. This method makes it possible to estimate not only the effects of the studied factors (concentrations of copper ions, densities, etc.) on mortality and rates of development in common toad embryos, but also to reveal the effects of interaction between factors and thus test more complex hypotheses. Therefore, factorial ANOVA can be used to estimate the interaction between factors described as changes in the effect of one factor that result from another factor. The normality hypotheses and homogeneity were analyzed with Lillieford and Kolmogorov-Smirnov tests, respectively, using Statistica 6.0 software.

## RESULTS

### Copper toxicity

Temperature conditions affect the rates of development and mortality in common toad embryos (Dmitrieva, 2014). The duration of embryonic development until hatching was 130 h in clutches that developed in a thermostat at 15°C (trial 2) and 230 h in clutches that developed in a thermostatic room at 10–13°C (trial 1). Thus, the conditions under which embryos developed in trial 1 and trial 2 were different, and therefore the trials were analyzed separately.

Short-term toxicity test (48 h) shows that mortality was almost zero in all clutches at a concentration of 10 mg/L and 25 mg/L (Table 1). At a high concentration of Cu ions (50 mg/L), embryo mortality increases (Table 1); it is significantly higher (by Kruskal-Wallis test) in clutch 3 than under normal conditions at a density of both 30 eggs per aquarium (*H*(3, *N* = 29) = 21.238, *p* = 0.0001) and 120 eggs per aquarium (*H*(3, *N* = 13) = 10.164, *p* = 0.0172). In trial 2 mortality was significantly higher (by Kruskal-Wallis test) than under normal conditions in clutch 4 only at higher density ((*H* (3, *N* = 12) = 8.214, *p* = 0.0418) and (*H*(3, *N* = 16) = 11.058, *p* = 0.0114), respectively) and, by contrast, only at lower density (*H*(3, *N* = 16) = 14.619, *p* = 0.0022) in clutch 5.

**Table 1.** Embryo mortality (% mean mortality ± SE) in short-term toxicity tests (48 and 96 h)

| Tests | Trials | Clutch | Density<sup>1</sup> | Cu concentration |  |  |  |
| --- | --- | --- | --- | --- | --- | --- | --- |
|  |  |  |  | 0 mg/L | 10 mg/L | 25 mg/L | 50 mg/L |
| 48 h | Trial 1 | 1 | 30 | 0.00±0.00 | 0.00±0.00 | 0.00±0.00 | 5.00±3.19 |
| | | | 120 | 0.00±0.00 | 0.00±0.00 | 0.00±0.00 | 3.13±1.38 |
| | | 2 | 30 | 0.00±0.00 | 0.00±0.00 | 0.00±0.00 | 11.67±2.15<sup>2</sup> |
| | | | 120 | 0.00±0.00 | 0.00±0.00 | 0.21±0.21 | 3.54±1.61 |
| | | 3 | 30 | 0.00±0.00 | 0.00±0.00 | 6.67±4.71 | 29.17±4.59<sup>2</sup> |
| | | | 120 | 0.00±0.00 | 0.00±0.00 | 0.56±0.56 | 5.83±0.96<sup>2</sup> |
| | Trial 2 | 4 | 30 | 0.83±1.67 | 0.00±0.00 | 1.67±1.92 | 4.17±3.19 |
| | | | 120 | 0.00±0.00 | 0.28±0.48 | 0.28±0.48 | 4.44±1.69<sup>2</sup> |
| | | 5 | 30 | 0.00±0.00 | 0.83±0.83 | 0.00±0.00 | 3.33±1.36 |
| | | | 120 | 0.56±0.28 | 0.00±0.00 | 0.42±0.24 | 5.00±0.76<sup>2</sup> |
| 96 h | Trial 1 | 1 | 30 | 0.00±0.00 | 0.00±0.00 | 1.67±1.92 | 19.17±2.50<sup>2</sup> |
| | | | 120 | 0.00±0.00 | 0.00±0.00 | 1.04±0.21<sup>2</sup> | 12.92±4.34<sup>2</sup> |
| | | 2 | 30 | 0.00±0.00 | 0.00±0.00 | 8.33±3.19 | 50.83±3.94<sup>2</sup> |
| | | | 120 | 0.00±0.00 | 0.63±0.63 | 7.08±2.49<sup>2</sup> | 40.00±6.26<sup>2</sup> |
| | | 3 | 30 | 0.00±0.00 | 0.00±0.00 | 19.17±13.77 | 94.17±2.10<sup>2</sup> |
| | | | 120 | 0.00±0.00 | 0.00±0.00 | 2.78±0.73<sup>2</sup> | 42.78±5.74<sup>2</sup> |
| | Trial 2 | 4 | 30 | 0.83±1.67 | 80.00±9.03<sup>2</sup> | 100.00±0.00<sup>2</sup> | 100.00±0.00<sup>2</sup> |
| | | | 120 | 0.56±0.48 | 52.78±8.01<sup>2</sup> | 46.11±4.74<sup>2</sup> | 58.06±5.19<sup>2</sup> |
| | | 5 | 30 | 2.50±0.83 | 90.83±5.99<sup>2</sup> | 97.50±1.60<sup>2</sup> | 100.00±0.00<sup>2</sup> |
| | | | 120 | 0.83±0.00 | 28.54±3.91<sup>2</sup> | 52.71±9.24<sup>2</sup> | 77.50±8.46<sup>2</sup> |

<sup>1</sup> initial density (number of eggs per aquarium)<sup>2</sup> indicates values of mortality significantly ( $p < 0.05$ ) different from control group according to Mann–Whitney test (Cu concentration 0 mg/L)

In the 96 h test in trial 2, mortality was significantly higher than under normal conditions at any studied concentration of Cu ions: *H*(3, *N* = 16) = 14.619, *p* = 0.0022 at a density of 30 eggs per aquarium in clutch 4; *H*(3, *N* = 12) = 8.379, *p* = 0.0388 at a density of 120 eggs per aquarium in clutch 4; *H*(3, *N* = 16) = 11.529, *p* = 0.0092 at a density of 30 eggs per aquarium in clutch 5; and *H* (3, *N*= 16) = 12.870, *p* = 0.0049 at a density of 120 eggs per aquarium in clutch 5. In trial 1, in which embryos developed at decreased temperature, mortality was significantly higher only at high concentrations of copper ions (50 mg/L), while at medium concentrations (25 mg/L) mortality was significantly higher than in the control group only at low egg density (30 eggs per aquarium) (Table 1).

The long-term test, which lasted until the hatching of larvae in the control group, shows that all studied concentrations of copper ions prove lethal for common toad embryos (Fig. 1), i.e., by the moment of hatching in the control group all eggs were dead in the experimental groups. Survival rate in the control group was 96–100%, independently of egg density or temperature at which the embryos developed. Embryos developed for approximately 230 h at 11–13°C and 130 h at 15°C. The timing of the end of mortality (the moment when the last embryo died) differed depending on developmental conditions (Table 2). For instance, in clutch 1 at a concentration of 50 mg/L mortality started approximately 40–48 h after the start of the experiment, and all embryos died approximately by hour 192–200 of the experiment. As for clutches 2 and 3, mortality in them started at the same time, but embryos died much earlier at a density of 30 eggs per aquarium. At 10 mg/L mortality began much later than at 50 mg/L (Table 2, Fig. 1). This difference was especially pronounced in clutches 1–3 that developed at lower temperatures: embryos began to die 104–112 h after the start of the experiment.

**Fig.1.**
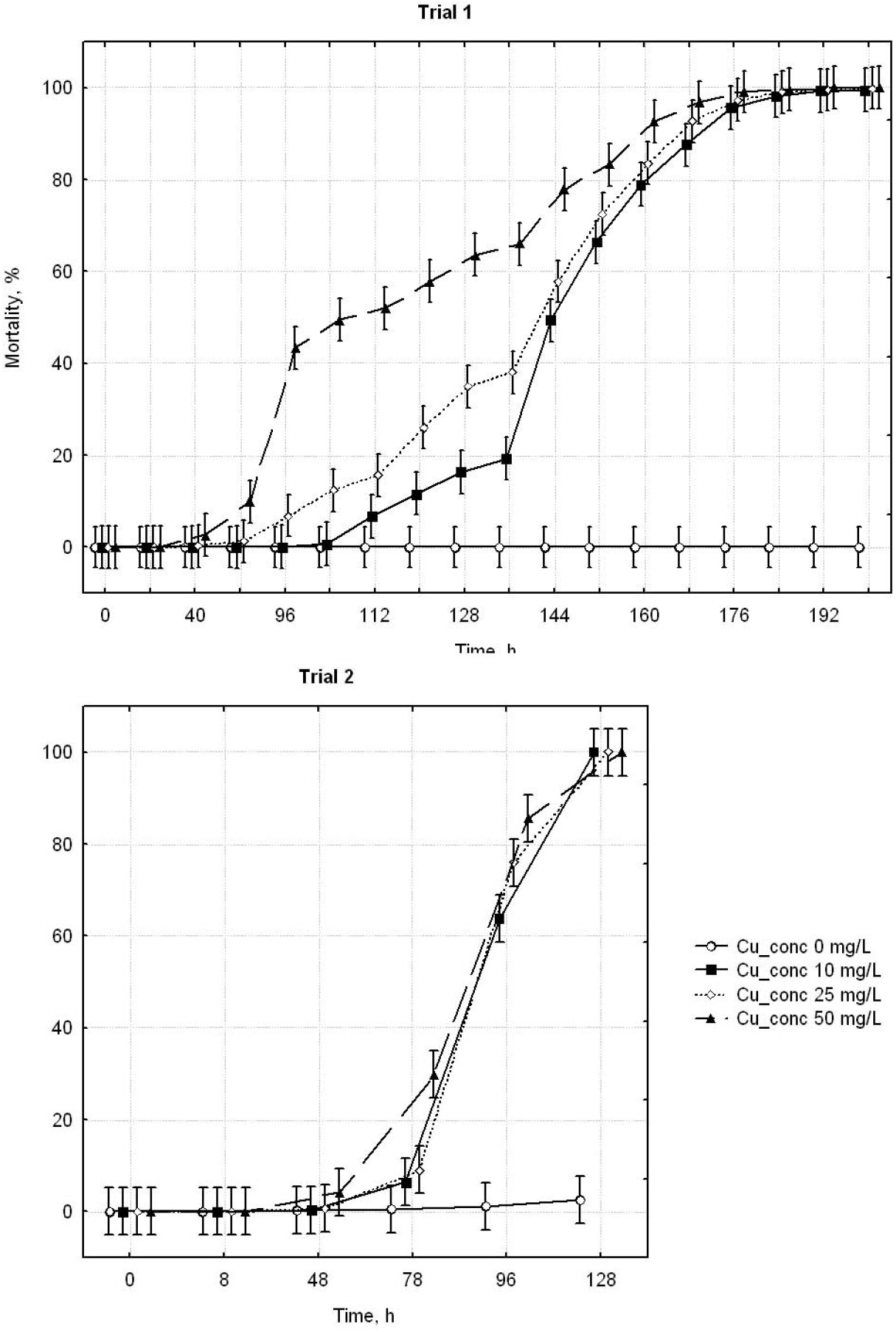
Effect of aqueous copper concentration on embryo mortality in Trial 1 (a) and in Trial 2 (b) in *Bufo bufo*. Treatment means (±1 SE) for each point are shown.

**Table 2.** Beginning and end of mortality (all individuals dead) in *Bufo bufo* embryos in different clutches at different initial densities and Cu concentrations

| Trials | Clutch | Density <sup>1</sup> | Beginning of mortality, h <sup>2</sup> |  |  |  | All embryos dead, h <sup>2</sup> |  |  |  |
| --- | --- | --- | --- | --- | --- | --- | --- | --- | --- | --- |
|  |  |  | Cu concentration |  |  |  | Cu concentration |  |  |  |
|  |  |  | 0 mg/L | 10 mg/L | 25 mg/L | 50 mg/L | 0 mg/L | 10 mg/L | 25 mg/L | 50 mg/L |
| Trial 1 | 1 | 30 | No <sup>3</sup> | 104 | 96 | 48 | No <sup>3</sup> | 192 | 176 | 192 |
|  |  | 120 | No <sup>3</sup> | 112 | 56 | 40 | No <sup>3</sup> | 200 | 184 | 192 |
|  | 2 | 30 | No <sup>3</sup> | 112 | 80 | 48 | No <sup>3</sup> | 184 | 192 | 176 |
|  |  | 120 | No <sup>3</sup> | 56 | 48 | 40 | No <sup>3</sup> | 232 | 200 | 192 |
|  | 3 | 30 | 223 | 104 | 48 | 40 | No <sup>3</sup> | 192 | 200 | 144 |
|  |  | 120 | No <sup>3</sup> | 104 | 48 | 40 | No <sup>3</sup> | 216 | 224 | 200 |
| Trial 2 | 4 | 30 | 78 | 78 | 48 | 48 | No <sup>3</sup> | 128 | 102 | 102 |
|  |  | 120 | 78 | 56 | 48 | 48 | No <sup>3</sup> | 128 | 128 | 128 |
|  | 5 | 30 | 102 | 56 | 56 | 48 | No <sup>3</sup> | 128 | 128 | 102 |
|  |  | 120 | 78 | 78 | 78 | 48 | No <sup>3</sup> | 128 | 128 | 128 |

<sup>1</sup> initial density (number of eggs per aquarium)
<sup>2</sup> hour since start of experiment
<sup>3</sup> not all tadpoles died

According to the Mann–Whitney test, in trial 1 at 50 mg/L mortality was significantly higher than at 10 mg/L from 96 to 168 h (p < 0.0005) and 176 h (*p* = 0.018) after the start of development, and significantly higher than at 25 mg/L from 96 to 144 h (*p* < 0.005) and 160 h (*p* < 0.005) after the start of development. Significant differences in mortality between 10 and 25 mg/L have been revealed from 96 to 136 h (*p* < 0.005) and 168 h (*p* = 0.016) after the start of development. Subsequently, the differences became insignificant again, and mortality in all series of the experiment approached 100%. In trial 2, mortality at 50 mg/L was significantly higher than at 10 mg/L from 48 to 102 h (*p* < 0.005) after the start of development and significantly higher than at 25 mg/L from 48 to 78 h (*p* < 0.005) after the start of development. Subsequently, the differences became insignificant again, and mortality in all series of the experiment approached 100%. Mortality at 10 and 25 mg/L did not differ significantly.

The influence of copper began to manifest itself at the blastula stage (Fig. 2a), when group mortality of embryos was first recorded. Copper ions slowed down embryonic development, terminating it at the mid-gastrula stage (Fig. 2b), while in the control group development continued, embryos freely developed, and eventually hatched as larvae (stage 19) (Gosner, 1960). Therefore, in the experimental groups development at all concentrations of copper ions was decelerated, compared to the control group, and embryonic development was terminated at the gastrula stage, followed by mass mortality. This pattern was recorded in all studied clutches, independently of temperature or density at which the embryos developed.

**Fig.2.**
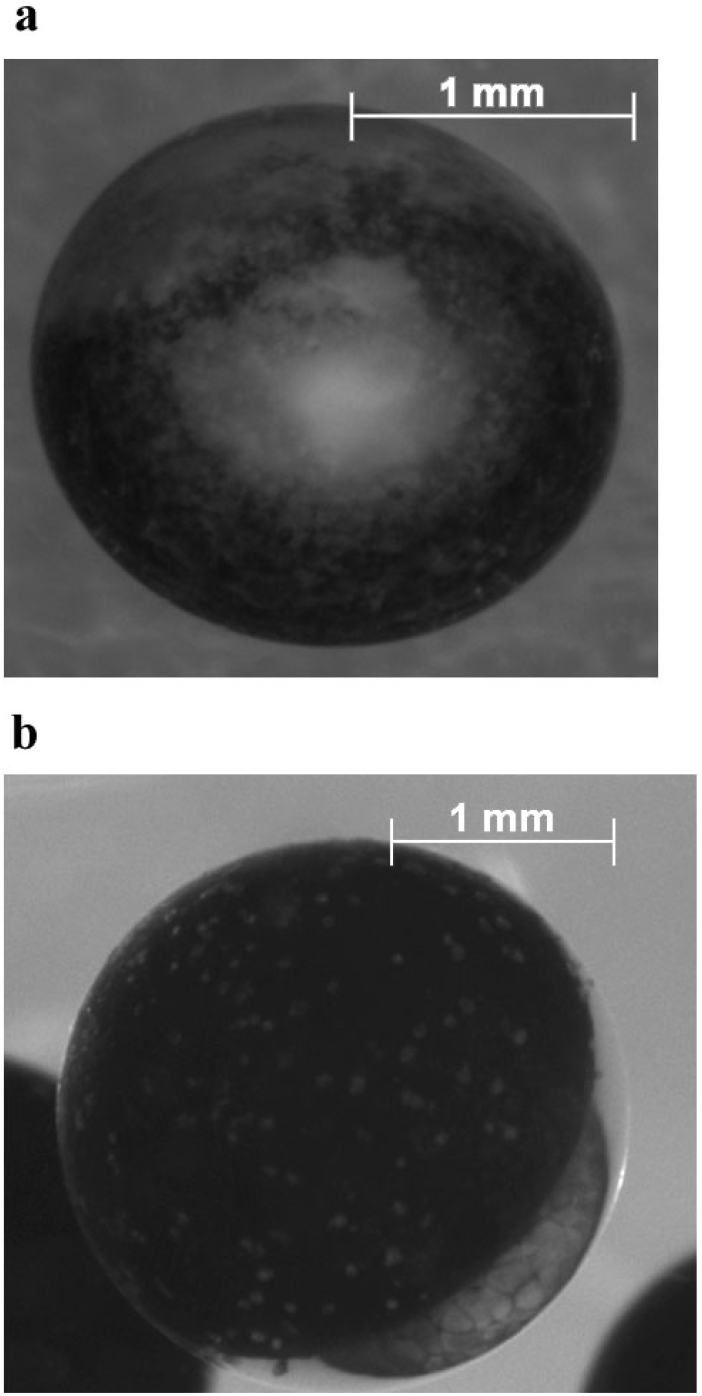
Optical microscopic views of malformations produced by Cu in Bufo bufo embryos in mid-blastula stage (a) and gastrula stage (b).

### Differences between clutches and effects of density

Factorial ANOVA shows significant influence of all studied factors on mortality of embryos in both 48 h test and 96 h test in clutches 1–3 (trial 1), which developed at a lower temperature (Table 3). In clutches 4–5 (trial 2) mortality was affected above all by the concentration of Cu ions (in both 48 h test and 96 h test). In the 96 h test, mortality was also affected by density of developing embryos, and one factor changed under the influence of the other (Table 3). In the short-term (48 h) test, mortality was affected only by the concentration of Cu ions in clutch 1 (trial 1), as well as in clutches 4 and 5 (trial 2), whereas in clutches 2 and 3 (trial 1) it was also affected by density of developing embryos (Table 4). In the 96 h test, mortality was affected by both factors in clutches 4 and 5 (trial 2), whereas in trial 1 density had a significant effect on mortality only in clutch 3 (Table 4).

**Table 3.** Results of factorial ANOVA: mortality as the dependent variable and copper concentration, clutches and density as the categorical variables

| Trial | Tests | Source of variation | *df* | MS | *F* | *p* |
| --- | --- | --- | --- | --- | --- | --- |
| Trial 1<br>(clutches<br>1-3) | 48 h | Clutch | 2 | 147.5878 | 13.22315 | 0.000014 |
|  |  | Density | 1 | 243.0746 | 21.77830 | 0.000015 |
|  |  | Cu concentration | 3 | 500.6881 | 44.85921 | 0.000000 |
|  |  | Clutch*Density | 2 | 96.1418 | 8.61384 | 0.000463 |
|  |  | Clutch*Copper concentration | 6 | 79.5607 | 7.12825 | 0.000006 |
|  |  | Density*Copper concentration | 3 | 160.3410 | 14.36577 | 0.000000 |
|  |  | Clutch*Density* Copper concentration | 6 | 50.2944 | 4.50613 | 0.000671 |
|  |  | Error | 68 | 11.1613 |  |  |
|  | 96 h | Clutch | 2 | 1801.97 | 33.1245 | 0.000000 |
|  |  | Density | 1 | 1170.81 | 21.5223 | 0.000016 |
|  |  | Cu concentration | 3 | 9807.65 | 180.2882 | 0.000000 |
|  |  | Clutch*Density | 2 | 526.94 | 9.6864 | 0.000199 |
|  |  | Clutch*Copper concentration | 6 | 1169.47 | 21.4977 | 0.000000 |
|  |  | Density*Copper concentration | 3 | 667.33 | 12.2672 | 0.000002 |
|  |  | Clutch*Density* Copper concentration | 6 | 249.26 | 4.5820 | 0.000583 |
|  |  | Error | 68 | 54.40 |  |  |
| Trial 2<br>(clutches<br>4-5) | 48 h | Clutch | 1 | 0.6410 | 0.26374 | 0.610134 |
|  |  | Density | 1 | 0.0000 | 0.00000 | 1.000000 |
|  |  | Cu concentration | 3 | 54.8077 | 22.54945 | 0.000000 |
|  |  | Clutch*Density | 1 | 2.5641 | 1.05495 | 0.309984 |
|  |  | Clutch*Copper concentration | 3 | 0.6766 | 0.27839 | 0.840685 |
|  |  | Density*Copper concentration | 3 | 1.6026 | 0.65934 | 0.581474 |
|  |  | Clutch*Density* Copper concentration | 3 | 1.6026 | 0.65934 | 0.581474 |
|  |  | Error | 44 | 2.4306 |  |  |
|  | 96 h | Clutch | 1 | 32.5 | 0.458 | 0.502057 |
|  |  | Density | 1 | 14981.3 | 210.893 | 0.000000 |
|  |  | Cu concentration | 3 | 20500.8 | 288.592 | 0.000000 |
|  |  | Clutch*Density | 1 | 15.2 | 0.214 | 0.645579 |
|  |  | Clutch*Copper concentration | 3 | 166.9 | 2.349 | 0.085475 |
|  |  | Density*Copper concentration | 3 | 1747.3 | 24.596 | 0.000000 |
|  |  | Clutch*Density* Copper concentration | 3 | 515.9 | 7.263 | 0.000463 |
|  |  | Error | 44 | 71.0 |  |  |

It should be emphasized one again that the timing of embryo mortality was different in different clutches. Thus, at the highest Cu concentration at a density of 30 eggs per aquarium all embryos died much sooner in clutches 2 and 3 than in clutch 1: by hour 176, 148, and 192, respectively (Table 2). In the long-term toxicity test, the end of mortality (the moment when the last embryo died) was timed differently in different clutches (Table 2). In clutches 2 and 3 at low density of embryos they all died much earlier: 176 and 148 h, respectively, after the start of the experiment. In addition, It should be noted that in almost all studied clutches embryos died earlier at a density of 30 eggs per aquarium than at 120 eggs per aquarium.

The clutches used in trial 1 differed in sensitivity to copper levels (Fig. 3). At early stages of ontogenesis, clutch 1 was more resistant to the presence of copper ions than clutches 2 and 3. According to the Mann–Whitney test, mortality in clutch 1 remained significantly lower until 160 h after the start of development at 10 mg/L (*p* < 0.05) and 50 mg/L (*p* < 0.05) and until 144 h after the start of development at 25 mg/L (*p* < 0.05). However, subsequently mortality rate in clutch 1 increased, and all individuals were dead much earlier than in the other clutches. Clutch 2 proved the most sensitive to lower concentrations of copper ions (10 mg/L and 25 mg/L): mortality was significantly (*p* < 0.05) lower than in clutches 1 and 3. However, at high concentrations of copper ions (50 mg/L) clutch 3 was the most sensitive. In clutches 4 and 5 (trial 2) mortality was almost identical.

**Fig.3.**
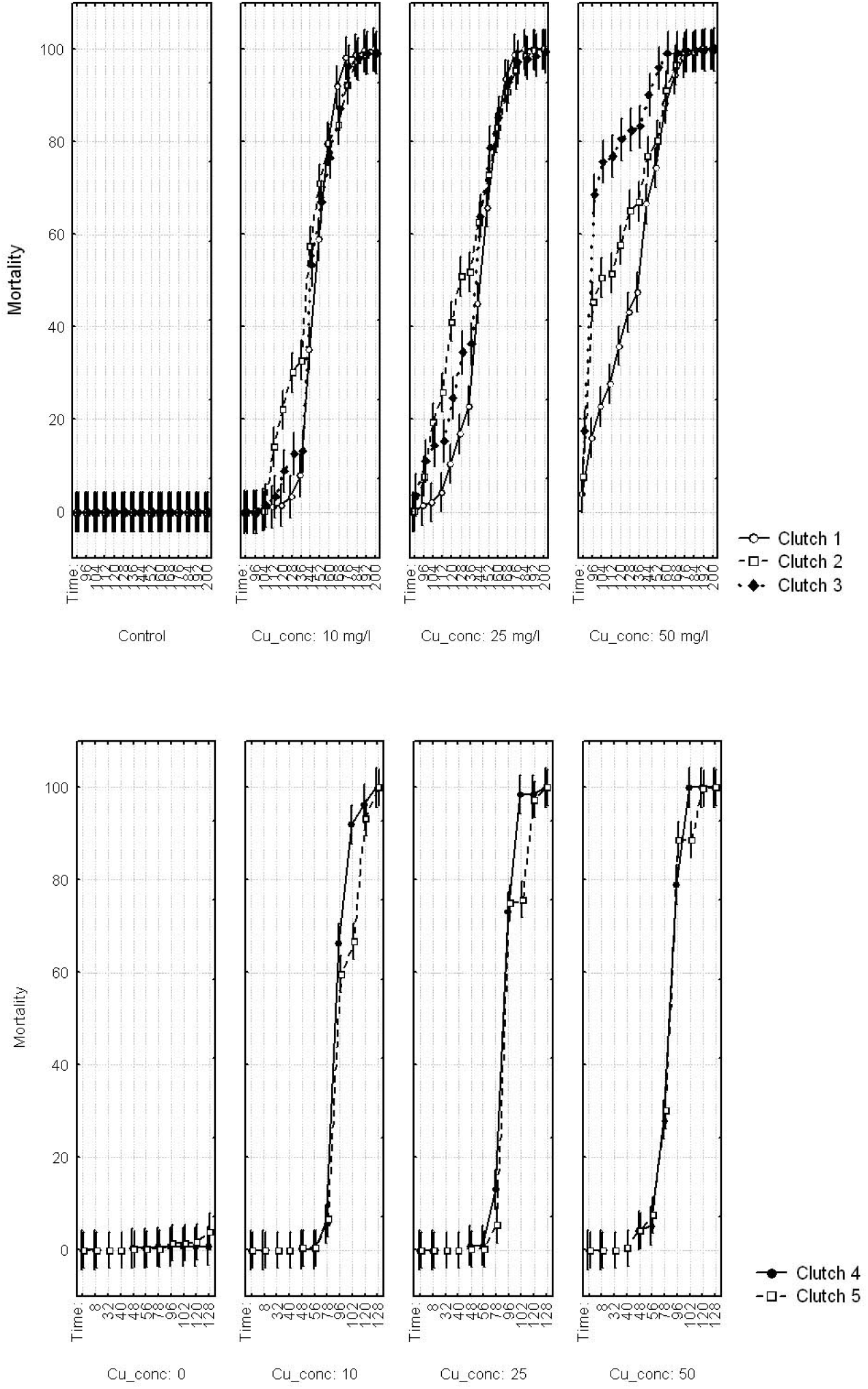
Effect of aqueous copper concentration on embryo mortality in different clutches. Treatment means (±1 SE) for each point are shown.

The density of developing embryos can also affect mortality at different copper levels (Table 3). In the short-term toxicity test (48 h), density and its interaction with Cu concentration affected two studied clutches in trial 1 (Table 4). The 96 h toxicity test revealed a similar effect in clutches 4 and 5 (trial 2) that developed at 15°C (Table 4), and in one other clutch that developed at lower temperatures (clutch 3; Table 4). In the long-term experiment, density affected mortality at different copper levels in all studied clutches (Table 5). Moreover, almost all studied factors affected the resulting mortality of embryos.

**Table 4.** Results of factorial ANOVA: mortality as the dependent variable and copper concentration and density as the categorical variables in 48-h and 96-h toxicity tests

| Clutches | Tests | Source of variation | df | MS | F | p |
| --- | --- | --- | --- | --- | --- | --- |
| Clutch 1 | 48 h | Density | 1 | 1.68750 | 0.267737 | 0.609795 |
|  |  | Cu concentration | 3 | 32.60031 | 5.172322 | 0.007045 |
|  |  | Density*Cu concentration | 3 | 1.73611 | 0.275449 | 0.842487 |
|  |  | Error | 23 | 6.30284 |  |  |
|  | 96 h | Density | 1 | 22.6875 | 1.67073 | 0.208996 |
|  |  | Cu concentration | 3 | 481.9016 | 35.48767 | 0.000000 |
|  |  | Density*Cu concentration | 3 | 18.1327 | 1.33531 | 0.287311 |
|  |  | Error | 23 | 13.5794 |  |  |
| Clutch 2 | 48 h | Density | 1 | 31.3368 | 8.62948 | 0.007194 |
|  |  | Cu concentration | 3 | 114.6123 | 31.56175 | 0.000000 |
|  |  | Density*Cu concentration | 3 | 33.5938 | 9.25100 | 0.000301 |
|  |  | Error | 24 | 3.6314 |  |  |
|  | 96 h | Density | 1 | 65.647 | 1.8359 | 0.188046 |
|  |  | Cu concentration | 3 | 3755.462 | 105.0283 | 0.000000 |
|  |  | Density*Cu concentration | 3 | 57.661 | 1.6126 | 0.212647 |
|  |  | Error | 24 | 35.757 |  |  |
| Clutch 3 | 48 h | Density | 1 | 385.3224 | 15.35872 | 0.000788 |
|  |  | Cu concentration | 3 | 485.3192 | 19.34453 | 0.000003 |
|  |  | Density*Cu concentration | 3 | 213.5520 | 8.51206 | 0.000682 |
|  |  | Error | 21 | 25.0882 |  |  |
|  | 96 h | Density | 1 | 2041.70 | 16.95561 | 0.000490 |
|  |  | Cu concentration | 3 | 7529.36 | 62.52869 | 0.000000 |
|  |  | Density*Cu concentration | 3 | 1032.51 | 8.57465 | 0.000655 |
|  |  | Error | 21 | 120.41 |  |  |
| Clutch 4 | 48 h | Density | 1 | 1.19048 | 0.34985 | 0.560822 |
|  |  | Cu concentration | 3 | 25.52910 | 7.50243 | 0.001487 |
|  |  | Density*Cu concentration | 3 | 1.19048 | 0.34985 | 0.789665 |
|  |  | Error | 20 | 3.40278 |  |  |
|  | 96 h | Density | 1 | 6519.05 | 221.750 | 0.000000 |
|  |  | Cu concentration | 3 | 9101.63 | 309.599 | 0.000000 |
|  |  | Density*Cu concentration | 3 | 915.26 | 31.133 | 0.000000 |
|  |  | Error | 20 | 29.40 |  |  |
| Clutch 5 | 48 h | Density | 1 | 1.38889 | 0.85714 | 0.363756 |
|  |  | Cu concentration | 3 | 30.32407 | 18.71429 | 0.000002 |
|  |  | Density*Cu concentration | 3 | 2.08333 | 1.28571 | 0.301923 |
|  |  | Error | 24 | 1.62037 |  |  |
|  | 96 h | Density | 1 | 8640.6 | 81.7188 | 0.000000 |
|  |  | Cu concentration | 3 | 11771.4 | 111.3276 | 0.000000 |
|  |  | Density*Cu concentration | 3 | 1384.0 | 13.0891 | 0.000029 |
|  |  | Error | 24 | 105.7 |  |  |

The lower was the concentration of copper ions, the later did density begin to affect mortality. For instance, according to the Mann–Whitney test, in trial 1 at 50 mg/L mortality at a density of 30 and 120 eggs per aquarium significantly differed from 48 to 56 h and from 112 to 152 h after the start of the experiment (*p* < 0.05), whereas at 10 mg/L it differed significantly from 136 to 200 h after the start of the experiment (*p* < 0.05). In trial 2 mortality at 30 and 120 eggs per aquarium significantly differed from 78 to 102 h after the start of the experiment at 10 mg/L (*p* < 0.002) and 25 mg/L (*p* < 0.05). At 50 mg/L mortality at a density of 30 and 120 eggs per aquarium significantly differed only from 96 to 102 h after the start (*p* < 0.006).

## DISCUSSION

The long-term toxicity test revealed the lethal effect of each of the studied concentrations of copper ions on common toad eggs: all individuals were dead in the experimental groups by the time of hatching in the control group. This was the expected result, since the investigated copper concentrations were certainly above the limits of amphibians tolerance to Cu (Herkovits, Helguero, 1998; Lance et al., 2012, 2013; Weir et al., 2016a). Under natural conditions such high levels of copper concentration were not observed. However, in spite of the high concentrations of copper in our experiments, the short-term (48 h) toxicity test revealed no significant differences in mortality at 10 mg/L and 25 mg/L. Mortality increased only at the highest tested concentration of copper ions (50 mg/L). The 96 h test revealed significantly increased mortality at all studied concentrations only in trial 2, at a higher temperature than in trial 1. In trial 1 mortality was significantly higher only at the highest concentration and in some cases at 25 mg/L. Therefore, short-term toxicity tests can fail to reveal the toxic effect of copper even at such high concentrations of copper ions as in the present study. Let us discuss in more detail why such results could be obtained.

The results of toxicity tests of the same substance can differ depending on the method. For instance, the AMPHITOX method (Herkovits, Pérez-Coll, 2003) is based on biotesting in ecosystem models (mesocosms). Ecotoxicological studies of the aquatic environment are performed by creating special experimental ponds with bottom sediments, macrophytes, mollusks, crustaceans, and fishes, which become stabilized over several months (Caquet et al., 1996). These mesocosms as such can partially utilize copper from the aquatic environment; as a result, the ecotoxicological effects of copper can differ from those revealed by other biotesting methods. Furthermore, high concentrations of other ions in the solution can compete with copper ions for binding and absorption, decreasing the bioaccumulation and toxicity of copper (Santore et al., 2001). FETAX and AMPHITOX tests differ also in the salinity and temperature of maintaining media (Caquet et al., 1996), which can also modify toxicity results.

According to the FETAX method (American Society for Testing and Materials, 1991) and some other studies (Edginton et al., 2004), it is recommended to remove the gelatinous egg membranes that protect developing embryos. The removal of these membranes results in increased mortality of amphibian embryos (Robinson et al., 2003), which can considerably influence the results of experiments. The same is true of the removal of dead embryos (Dmitrieva, 2005). In addition, the jelly coats of eggs can decrease the toxic effects of the environment at embryonic stages, prior to hatching (Aronzon et al., 2011). Therefore, in the present study a whole piece of the clutch, containing a particular number of eggs (30 or 120) was used in each experiment. Dead embryos were not removed, so as not to affect the results of the experiments: otherwise mortality could increase because of the influence of the experimenter rather than decrease, as might be expected (Dmitrieva, 2005).

In addition, the results of the experiment depend on the developmental stages affected by the exposure (Dregolskaya, 1993; Aronzon, 2011; Pérez-Coll C.S., Herkovits, 1990; Sztrum et al., 2011; Flynn et al., 2015). If testing is started at the cleavage stage (stage 2–4) (Gosner, 1960), the effects of copper are almost invisible. Short-term toxicity tests reveal only the effects of the highest copper levels. The early stages of development proved rather resistant to copper, even at high concentrations. During cleavage the influence of Cu on processes of survival and development remains invisible. At the mid-blastula stage (stage 8) (Gosner, 1960), increased embryo mortality is observed. At temperatures of 10–13°C (trial 1), the effects of low concentrations of copper on embryo mortality began to manifest themselves 104–112 h after the start of development, whereas at higher concentrations the effects were visible as early as 40–48 after the start of development (Table 2). At 15°C (trial 2) this difference became visible as early as 48 h after the start of development at each studied copper level, except for the lowest. The effects of copper ions on embryo mortality began to manifest themselves at the blastula stage (Fig. 2a), although the embryos continued to develop for some time until the mid-gastrula stage (Fig. 2b), at which development terminated. Previous studies on stage dependent susceptibility to noxious agents, including Cu, point out the high resistance of blastula stage amphibian embryos to this heavy metal (Xia et al., 2012). Therefore, under unfavorable conditions (increased copper level) embryonic development freezes at the gastrula stage, without switching to the stage of neural plate formation, as if waiting for unfavorable conditions to change for the better. If conditions do not improve, mass mortality begins, and can last up to several hours. A similar pattern of mortality was observed in *Rhinella arenarum* by Aronzon et al. (2011): development also stopped at the mid-gastrula stage (stage 11) (Gosner, 1960), and mortality resulted in cellular dissociation and yolk exudation to the perivitelline space, as in the present study (Fig. 2).

At the gastrulation stage, the rate of energy metabolism increases, increasing the consumption of microelements, including copper. However, if copper concentration in the environment is to high, copper can switch roles from activator of biological processes to toxicant. Outside their coordination spheres, copper ions become highly toxic agents, because they can catalyze processes of the sort of the Fenton reaction, resulting in production of active oxygen metabolites, which produces effects on all structures of the cell similar to the effects of radiation poisoning (Yoshida et al., 1993). Cell dissociation observed at the gastrula stage probably reflects the interaction of Cu and Ca, which results in destruction of intercellular contacts (Grosell, Wood, 2002). In addition, in long-term tests amphibian eggs can accumulate heavy metals, including copper (O’Dell, 1990), which can also affect the mortality of embryos.

Therefore, short-term toxicity tests performed at embryonic stages can fail to reveal the toxic effect of the substance in question because of the high stability of earlier stages of development (stages 2–8 (Gosner, 1960)). This can lead to erroneous predictions because amphibian susceptibility to Cu and other metals is highly stage dependent, with organogenic stages being the most susceptible and early embryonic and late larval stages the most tolerant (Aronzon et al., 2011; Pérez-Coll C.S., Herkovits, 1990; Sztrum et al., 2011). Thus, during the ecotoxicological investigations must take into account such factors as life stages affected by the Cu exposure.

Particular attention should be paid to the effects of density on common toad embryo mortality at different concentrations of copper. Planning experiments with embryonic stages of development, researchers pay little attention to the effects of egg density on the results of these experiments. Egg density is standardized only in the FETAX method (American Society for Testing and Materials, 1991). Although many researchers maintain the same density of embryos in replications of the same experiment, the parameter is not standardized even in studies of the same species. For instance, egg density in different experiments can vary from 20 per 10 mL of medium (Edginton et al., 2004) to 14–20 per 1.75 L of medium (Snodgrass et al., 2008), which makes interpretation and comparative analysis of the results of such studies extremely difficult. There are but few studies in which attention is paid to the method of biotesting and to taking into account the effects of density on the results of experiments. For instance, a study of the effects of copper concentration on mortality and development of embryonic and larval stages of *Lithobates (Rana) sphenocephalus* has shown that the results of the experiments can depend on initial egg density (Lance et al., 2013), a factor disregarded by most researchers planning such experiments.

The results of my experiments on the effects of copper have shown that density has a significant influence on embryo mortality in this species at different concentrations of copper ions (Table 5). I have shown earlier that the density of developing eggs has a considerable influence on the rate of development and embryo mortality in the common toad (Dmitrieva, 2005, 2007, 2013); moreover, not only the embryo density has an effect (number of embryos per unit of water volume), but also the geometry and position of the clutch in the aquarium (24). Female common toads lay their eggs in strings containing helically arranged eggs. The folding of this egg string can affect the mortality and development of embryos (Dmitrieva, 2007, 2013). In addition, it is important to take into account the effects of “linear” density (ratio of the number of eggs to the length of the strained egg string) and “surface” density (ratio of the number of eggs in an aquarium to the bottom area of the aquarium) (Dmitrieva, 2013). In the present study all these factors have been taken into account. Egg strings with either 30 or 120 eggs were all straightened and evenly distributed on the bottom of the aquarium; therefore, the results of the experiments could not be affected by geometry of the clutch at increased egg density.

**Table 5.** Results of factorial ANOVA: mortality as the dependent variable and copper concentration, clutches, time of development (i.e. hour since start of experiment) and density as the categorical variables in long-time toxicity test

| Trial | Clutches | Source of variation | df | MS | F | p |
| --- | --- | --- | --- | --- | --- | --- |
| Trial 1 | Clutch 1 | Density | 1 | 1275.6 | 36.76 | 0.000000 |
|  |  | Cu concentration | 3 | 86353.7 | 2488.34 | 0.000000 |
|  |  | Time of development | 14 | 28518.2 | 821.77 | 0.000000 |
|  |  | Density*Cu concentration | 3 | 482.3 | 13.90 | 0.000000 |
|  |  | Density Time | 14 | 106.2 | 3.06 | 0.000172 |
|  |  | Cu concentration*Time | 42 | 3544.9 | 102.15 | 0.000000 |
|  |  | Density* Cu concentration*Time | 42 | 53.4 | 1.54 | 0.021202 |
|  |  | Error | 345 | 34.7 |  |  |
|  | Clutch 2 | Density | 1 | 12176 | 359.02 | 0.000000 |
|  |  | Cu concentration | 3 | 122311 | 3606.57 | 0.000000 |
|  |  | Time of development | 14 | 19989 | 589.40 | 0.000000 |
|  |  | Density*Cu concentration | 3 | 1516 | 44.69 | 0.000000 |
|  |  | Density Time | 14 | 640 | 18.87 | 0.000000 |
|  |  | Cu concentration*Time | 42 | 2654 | 78.27 | 0.000000 |
|  |  | Density* Cu concentration*Time | 42 | 123 | 3.61 | 0.000000 |
|  |  | Error | 360 | 34 |  |  |
|  | Clutch 3 | Density | 1 | 13522.2 | 310.83 | 0.000000 |
|  |  | Cu concentration | 3 | 139956.2 | 3217.10 | 0.000000 |
|  |  | Time of development | 14 | 16937.3 | 389.33 | 0.000000 |
|  |  | Density*Cu concentration | 3 | 2104.1 | 48.37 | 0.000000 |
|  |  | Density Time | 14 | 551.3 | 12.67 | 0.000000 |
|  |  | Cu concentration*Time | 42 | 3265.9 | 75.07 | 0.000000 |
|  |  | Density* Cu concentration*Time | 42 | 197.6 | 4.54 | 0.000000 |
|  |  | Error | 315 | 43.5 |  |  |
| Trial 2 | Clutch 4 | Density | 1 | 2455.6 | 140.225 | 0.000000 |
|  |  | Cu concentration | 3 | 15539.4 | 887.382 | 0.000000 |
|  |  | Time of development | 3 | 33497.5 | 1912.877 | 0.000000 |
|  |  | Density*Cu concentration | 3 | 308.9 | 17.639 | 0.000000 |
|  |  | Density Time | 3 | 1454.4 | 83.052 | 0.000000 |
|  |  | Cu concentration*Time | 9 | 3810.3 | 217.589 | 0.000000 |
|  |  | Density* Cu concentration*Time | 9 | 234.2 | 13.373 | 0.000000 |
|  |  | Error | 80 | 17.5 |  |  |
|  | Clutch 5 | Density | 1 | 2271.1 | 24.546 | 0.000003 |
|  |  | Cu concentration | 3 | 18088.0 | 195.491 | 0.000000 |
|  |  | Time of development | 3 | 41199.9 | 445.281 | 0.000000 |
|  |  | Density*Cu concentration | 3 | 620.6 | 6.708 | 0.000370 |
|  |  | Density Time | 3 | 2136.3 | 23.088 | 0.000000 |
|  |  | Cu concentration*Time | 9 | 4514.0 | 48.787 | 0.000000 |
|  |  | Density* Cu concentration*Time | 9 | 308.0 | 3.329 | 0.001401 |
|  |  | Error | 1 | 2271.1 | 24.546 | 0.000003 |

Taking into account the joint action of egg density and the studied factor (in the present study, copper concentration) increases the precision of estimations of the effects of this factor and precision of the results obtained. Increased egg density in some cases results in decreased mortality from copper as a toxic agent. Similar results, with the toxic effect of copper decreasing with increasing egg density, have been obtained for *Lithobates sphenocephalus* (Lance et al., 2013). This decrease is probably explained by the protective role played by jelly coats of the eggs. The jelly coats bind copper ions; the higher is the density of eggs, the more dissolved copper will be bound, decreasing the level of Cu in the solution. Greater total area of jelly coats covering greater amounts of eggs results in binding of greater amounts of dissolved copper (Lance et al., 2013).

In the present study I have also analyzed the variation of responses not only between embryos but also between different clutches of eggs. This approach is rather rare in ecotoxicological studies of amphibians. Such studies usually include mixing of eggs or larvae from several clutches for uniform distribution of potential genetic effects. As a result, genetically uneven samples of individuals are obtained, and much information is lost. In the present study I have observed considerable variation between clutches in trial 1. Clutches differed in mortality and sensitivity to different concentrations of copper ions. I have also observed differences in several important temporal characteristics of the clutches, such as the beginning and end of mortality at different copper levels. These results are in agreement with those obtained for *Lithobates sphenocephalus* (Bridges, Semlitsch, 2001; Lance et al., 2013) and *Gastrophryne carolinensis* (Flynn et al., 2015). Considerable interclutch variation has been revealed in sensitivity to carbaryl (Bridges, Semlitsch, 2001) and copper (Lance et al., 2013; Flynn et al., 2015). This variation was explained by the additive component of genetic diversity. High interclutch variation in studies of sensitivity to these two very different pollutants has led researchers to suggestion that mixed eggs from a small number of clutches would not be adequately representative of the range of variation present in most populations. In my opinion, studies of uneven genetic material can result in inadequate estimations of the effects of the factor in question. If we study eggs taken from the same clutch produced by one pair of breeders, i.e., a clutch of full siblings, the contribution of the genetic component to individual variation is strongly decreased. Studying the responses of particular clutches, we can estimate interclutch variation, and comparing clutches produced by different populations, we can estimate interpopulation variation. This approach will help us estimate more precisely the possible toxic effects of the studied pollutant and its influence on animals at the individual, population, or species level. Assessment of tolerance to anthropogenic stress factors, such as ecological pollutants, and understanding the behavior of individuals, clutches, or populations under ecological stress are important for conservation biology studies and can improve estimations of the relations between amphibian population decline and environmental pollutants.

Thus, my results demonstrate clearly that exposure induces mortality, but that the effects can be clutch specific, and density and time dependent. In ecotoxicological experiments the experimenter should take into account such factors as the initial egg density and the geometry of egg distribution in an aquarium. In addition, individual sensitivity of clutches to the effects of a toxic agent can also vary, potentially affecting the results of the experiment. These considerations are especially important to limiting false negative results. Considering the results of the present study, in further studies I plan to analyze the effects of lower concentrations of copper (0.1 mg/L and less) on mortality and rates of development in early ontogenesis of *Bufo bufo*.

## ACKNOWLEDGEMENT

The present study was supported by the Russian Foundation for Basic Research (project no. 16-04-01771). I thank Petr N. Petrov for his assistance with the paper translation.

